# New whole genome *de novo* assemblies of three divergent strains of rice (*O. sativa*) documents novel gene space of *aus* and *indica*

**DOI:** 10.1101/003764

**Authors:** M.C. Schatz, L.G. Maron, J.C. Stein, Wences A. Hernandez, J. Gurtowski, E. Biggers, H. Lee, M. Kramer, E. Antoniou, E. Ghiban, M.H. Wright, J.H. Chia, D. Ware, S.R. McCouch, W.R. McCombie

## Abstract

The use of high throughput genome-sequencing technologies has uncovered a large extent of structural variation in eukaryotic genomes that makes important contributions to genomic diversity and phenotypic variation. Currently, when the genomes of different strains of a given organism are compared, whole genome resequencing data are aligned to an established reference sequence. However when the reference differs in significant structural ways from the individuals under study, the analysis is often incomplete or inaccurate. Here, we use rice as a model to explore the extent of structural variation among strains adapted to different ecologies and geographies, and show that this variation can be significant, often matching or exceeding the variation present in closely related human populations or other mammals. We demonstrate how improvements in sequencing and assembly technology allow rapid and inexpensive *de novo* assembly of next generation sequence data into high-quality assemblies that can be directly compared to provide an unbiased assessment. Using this approach, we are able to accurately assess the “pan-genome” of three divergent rice varieties and document several megabases of each genome absent in the other two. Many of the genome-specific loci are annotated to contain genes, reflecting the potential for new biological properties that would be missed by standard resequencing approaches. We further provide a detailed analysis of several loci associated with agriculturally important traits, illustrating the utility of our approach for biological discovery. All of the data and software are openly available to support further breeding and functional studies of rice and other species.

## Introduction

Rice (*Oryza sativa)* provides 20% of the world’s dietary energy supply and is the predominant staple food for 17 countries in Asia, 9 countries in North and South America and 8 countries in Africa. Within *O. sativa*, there are two major varietal groups, *Indica* and *Japonica*, that can be further subdivided into five major subpopulations: *indica* and *aus* share ancestry within the *Indica* varietal group, and *tropical japonica, temperate japonica* and *aromatic (Group V)* share ancestry within the *Japonica* varietal group (**Figure 1**) (1–3). The subpopulation structure of *O. sativa* is deep and ancient, with estimates of divergence showing average pairwise Fst values of 0.375 - 0.45 (1–3), compared to Fst values of 0.25 for dogs (4), around 0.10 to 0.12 across human populations (5), or 0.08-0.09 for heterotic groups in maize (6).

**Figure 1.**
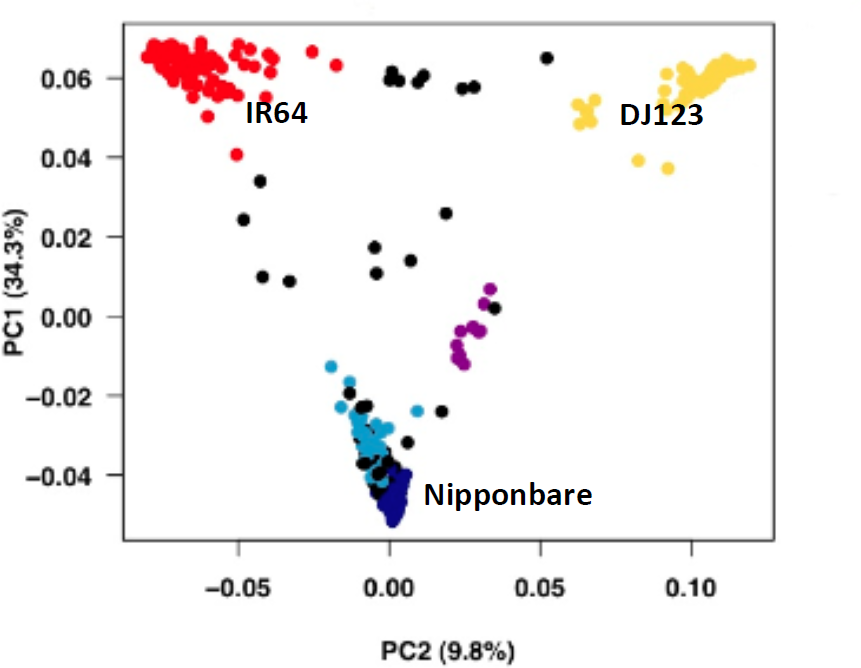
Population structure in *O. sativa.* A principal component analysis (PCA) based on 40,000 SNPs shows the deep subpopulation structure of a rice diversity panel (400 *O. sativa* accessions). The top two principal components (PCs) explain 44.1% of the genetic variation. Accessions are color-coded based on subpopulation: red, *indica*; dark blue, *temperate japonica*; light blue, *tropical japonica*; yellow, *aus;* purple, *aromatic;* black, admixed. Figure reproduced with permission from (83).

The time since divergence of the ancestral *Indica* and *Japonica* gene pools is estimated at 0.44 million years, based on sequence comparisons between cv Nipponbare (*Japonica)* and cv 93-11 (*Indica)* (7). This time estimate pre-dates the domestication of *O. sativa* by several hundred thousand years, suggesting that rice cultivation proceeded from multiple, pre-differentiated ancestral pools (1, 8–12). This is consistent with genome-wide estimates of divergence based on gene content (13), transcript levels (14), single nucleotide polymorphisms (SNPs) (3, 15), and transposable elements (16). This is also consistent with evidence from the cloning of dozens of genes underlying diverse quantitative trait loci (QTL) (2, 9, 17–21). Despite ongoing debate about the precise moment and location of the first domestication “event” in rice, these studies all demonstrate that natural variation in the rice genome is deeply partitioned and that divergent haplotypes can be readily associated with major varietal groups and subpopulations. The course of domestication, as rice transitioned from its ancestral state as a tropical, outcrossing, aquatic perennial species to a predominantly inbreeding, annual species adapted to a wide range of ecologies, was punctuated by persistent episodes of intermating among the different subpopulations. This resulted in both natural and human-directed gene flow between the different gene pools, but the essential differentiation that distinguishes the *Indica* and *Japonica* genomes was maintained and reinforced over time as a result of numerous partial sterility barriers scattered throughout the genome (22–25).

A better understanding of the nature and extent of genome variation within the *Oryza* clade is critical for both practical and scientific reasons. While the OMAP project (26) is focused on documenting structural variation across 21 wild species of *Oryza*, relatively little effort has been made to explore the nature of structural variation within and between subpopulations of *O. sativa.* The high quality, BAC-by-BAC sequence of the *temperate japonica* rice variety Nipponbare, generated by the International Rice Genome Sequencing Program (27), and the shotgun assembly of an *indica* rice genome, cv 93-11, by Chinese scientists in 2005 (28, 29) have served as ‘reference genomes’ for the rice research community. The availability of these reference genomes helped catalyze and unify rice research efforts for over a decade, and continue to serve as the backbone for re-sequencing efforts today (2, 30–33).

Recently, the resequencing of hundreds of wild and cultivated rice genomes using next generation sequencing (NGS) and various complexity-reduction and genotype-by-sequencing strategies have enriched the pool of sequence information available for rice (30, 34, 35). However, the vast majority of resequenced genomes are aligned to and compared with the Nipponbare reference rather than being assembled *de novo.* This introduces a potential bias due to significant differences in genome size (36, 37) and structure (13, 16, 29, 38) that characterize the different subpopulations and varieties of rice. Alignment to a single reference is particularly problematic when NGS from *indica, aus* or divergent wild species genomes from the center of diversity of *Oryza* are aligned to the genetically and geographically divergent Nipponbare (*temperate japonica)* reference because of the potential for misalignment, and for elimination of critical sequences that cannot be aligned with confidence.

The type and distribution of structural variation that distinguishes one rice genome from another, both within and between the five subpopulations of *O. sativa*, remains largely unknown. Yet it is essential to understanding the genetic basis of heterosis, as well as to identify genes underlying many of the most significant phenotypic differences that are critical to global food security, including a plant’s ability to grow in stressful environments afflicted by drought, submergence, low phosphorus and/or disease. The only practical way to fully understand the genomic diversity of rice is to carry out whole genome shotgun sequencing and *de novo* assembly. This has been problematic until recently due to the difficulties in assembling the short reads initially provided by NGS. However, recent advances in NGS chemistry and in computational approaches to sequence assembly have significantly improved the power and reliability of *de novo* assembly of NGS data.

In this study we use these new tools to develop *de novo* assemblies of three divergent rice genomes representing the *indica* (IR64), *aus* (DJ123) and *temperate japonica* (Nipponbare) subpopulations and to determine the extent and distribution of structural variation among them. These varieties were chosen for both biological interest and to facilitate evaluation of assemblies. On the biological side, different subpopulations of rice are adapted to different ecologies and geographies, and harbor different alleles and traits of interest for plant improvement (3, 18, 19, 39–42). The *aus* subpopulation is of particular interest because it is the source of important alleles conferring disease resistance (43), tolerance to submergence (33), deep water (44), low-phosphorus soils (40), and drought (45). *Indica* rice harbors the greatest amount of genetic variation (1, 30) and accounts for the largest contribution to rice production globally. Our choice to sequence Nipponbare was due to the fact that it provided a high quality BAC-by-BAC sequence assembly (27) that served as a solid benchmark for assessing the quality of our three NGS assemblies and provided a context for understanding the impact of varying data sets and parameters used in the assemblies.

## Results

### De novo genome assemblies and functional annotation

The three rice varieties were assembled using the ALLPATHS-LG whole genome assembler (46) using ∼50x coverage of a 180 bp fragment library, ∼30x coverage of a 2 kbp jumping library, and ∼30x coverage of a 5 kbp jumping library (see Methods). We selected this assembler based on its performance with these data compared to other assemblers and it’s high ranking in the Assemblathon I & II and GAGE evaluations (47–49). The three assemblies were named Os-Nipponbare-Draft-CSHL-1.0, Os-IR64-Draft-CSHL-1.0, and Os-DJ123-Draft-CSHL-1.0, following nomenclature proposed by (50).

All three of our assemblies had excellent results: approximately 90% of each of the genomes were assembled into scaffolds at least 1 kbp long, with scaffold N50 sizes ranging from 213 kbp to 323 kbp, and contig N50 sizes ranging from 21.9 kbp to 25.5 kbp (Table 1). It is notable that an earlier assembly of the Nipponbare genome prior to sequencing the 5 kbp jumping library achieved a similar contig N50 size (21.2 kbp versus 21.9 kbp), but a substantially smaller scaffold N50 size (99 kbp versus 213 kbp) (also see Methods). Improved scaffold sizes from including the larger library were expected, although the magnitude depends on the specific genome characteristics. Since the scaffolds were more than twice as large for Nipponbare with the larger library, this prompted us to sequence the 5 kbp jumping library for all three genomes to maximize our ability to identify genes and other features, as well as to structurally compare the genomes.

**Table 1.** Assembly and annotation statistics of the three de novo assemblies used in this study.

|  | **Nipponbare** | **IR64** | **DJ123** |
| --- | --- | --- | --- |
| **Total span** | 355.6 Mbp | 345.2 Mbp | 345.9 Mbp |
| **Total bases** | 318.2 Mbp | 316.3 Mbp | 321.2 Mbp |
| **Genome Coverage\***<br>(span / bases) | 91.2 % / 81.8 % | 88.5 % / 81.3 % | 88.6 % / 82.5 % |
| **Number of scaffolds** | 4,110 | 2,919 | 2,819 |
| **N50 scaffold span** | 213 kbp | 293 kbp | 323 kbp |
| **Max scaffold span** | 1.37 Mbp | 2.85 Mbp | 2.38 Mbp |
| **Number of contigs**<br>(>1000 bp) | 27,486 | 26,160 | 23,902 |
| **N50 contig size** | 21.9kbp | 22.2 kbp | 25.5 kbp |
| **Max contig size** | 133 kbp | 160 kbp | 252 kbp |
| **Total Genes** | 39,083 | 37,758 | 37,812 |
| **Median Gene Length** | 2,224 | 2,275 | 2,285 |
| **Mean exons per transcript** | 4.8 | 4.8 | 4.9 |

\*Assumes the total genome size is 389 Mbp, as according to IRGSP, 2005

The assemblies were repeat-masked and annotated for protein-coding genes using the MAKER-P automated pipeline (51), combining both evidence-based and *ab initio* methods (Supplementary Table 1). In addition to EST and full-length cDNA, we included as evidence the two published annotations of Nipponbare (50), and the published annotations of strains 93-11 and PA64s (28), thereby maximizing consistency and reducing bias of annotation across the three assemblies. Putative transposon-encoded genes were screened following analysis of InterPro domains (see Methods), which flagged ∼1% of initial gene calls in each of the three assemblies. Summary statistics for remaining genes are provided in Table 1 and Supplementary Table 2. Gene counts ranged from 37,758 (IR64) to 39,083 (Nipponbare), similar to the numbers reported by the MSU and RAP projects for the Os-Nipponbare-Reference-IRGSP-1.0 (39,102 and 35,681 respectively) (50). Overall statistics for structural features, such as exons, introns, and coding regions were highly consistent between the three assemblies and with published annotations. For instance, average translated protein lengths compared across MSU, RAP, and the three *de novo* assemblies ranged from 280-288 aa (median values: 268-291 aa), suggesting that contiguity of the *de novo* assemblies did not limit ability to identify protein-coding genes. For each assembly, 61-62% of annotated loci possessed one or more InterPro domain and 77% showed homology to plant NCBI RefSeq genes.

### Whole Genome Comparison to Nipponbare reference genomes

We evaluated the agreement between our *de novo* assemblies to the Nipponbare reference sequences using the GAGE assembly evaluation algorithm (49). As expected, the *de novo* Nipponbare assembly very closely matches the reference Nipponbare sequence, with a 99.94% average percent identity and only 0.31% of the assembly not aligning to the reference (Table 2). Even at this very high agreement, there are several tens of thousands of small variations, and several hundred larger variations. These variations are a combination of true variations from our sample relative to the reference genome, of which we expect there to be few, and errors from ALLPATHS-LG when used with these libraries and coverage levels. Consequently, considering that the assembly has a 99.94% overall similarity, the upper-bound on the error rate of sequencing and assembling with ALLPATHS-LG is at most 0.06%.

**Table 2a.** Comparison of the three de novo assemblies to the Nipponbare reference (IRGSP-1.0)

|  | **Unaligned**<br>**Reference**<br>**Bases** | **Unaligned**<br>**Assembly**<br>**Bases** | **Avg ID** | **SNPs &**<br>**Small**<br>**Indels** | **Indels**<br>**>5bp** | **Inver.** | **Reloc.** | **Trans.** |
| --- | --- | --- | --- | --- | --- | --- | --- | --- |
| Nipponbare | 3.14% | 0.31% | 99.94% | 57,459 | 3,445 | 131 | 252 | 617 |
| IR64 | 11.08% | 8.94% | 98.91% | 2,917,780 | 80,631 | 1,004 | 1,721 | 7,060 |
| DJ123 | 9.85% | 8.55% | 98.93% | 2,933,257 | 80,346 | 1,007 | 1,615 | 6,683 |

**Table 2b.**
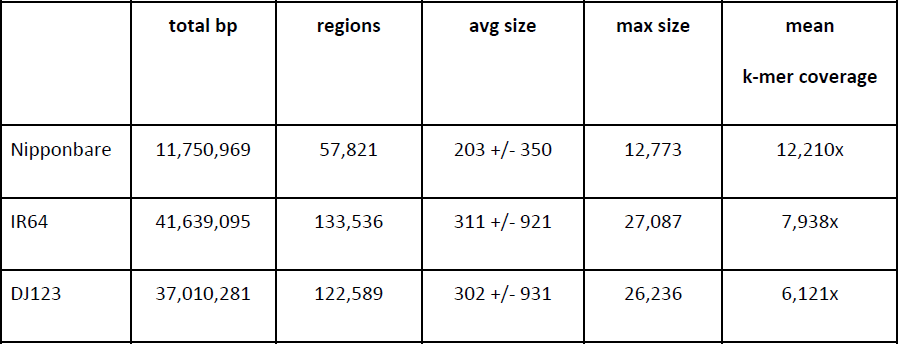
Summary of unaligned reference regions. Mean k-mer coverage was evaluated by counting the k-mers in a sample of 400M unassembled reads in each of the three genomes, and evaluating those counts along the reference sequence

**Table 2c.** Summary of unaligned reference regions. Mean k-mer coverage was evaluated by counting the k-mers in a sample of 400M unassembled reads in each of the three genomes, and evaluating those counts along the reference sequence

|  | **non-coding**<br>**exon** | **5' UTR** | **3' UTR** | **mRNA** | **CDS** | **repeat bp (>100x**<br>**k-mer coverage)** |
| --- | --- | --- | --- | --- | --- | --- |
| Nipponbare | 12,344 | 75,443 | 80,518 | 984,489 | 301,525 | 10,863,120 |
| IR64 | 129,428 | 902,520 | 638,856 | 6,827,679 | 1,706,454 | 29,685,919 |
| DJ123 | 114,735 | 865,470 | 601,928 | 6,418,865 | 1,519,686 | 24,505,588 |

The portions of the reference genome without any alignments from our Nipponbare assembly are scattered throughout the genome in 57,821 segments averaging 203 bp long. However, of this only 301,525 bp are annotated to be within the CDS (0.72% of the total CDS), and another 12,344 bp are annotated to be within non-coding exons. We further evaluated the unaligned regions by computing their read k-mer coverage from a sample of 400 million unassembled Nipponbare reads, and found the mean k-mer coverage of these regions exceeds 12,000x, while the mode k-mer coverage of the set is less than 100x (Supplementary Figure 1). A full two-thirds (38,373/57,821) of these regions exceed 1,000x k-mer coverage, more than 10 times higher than unique segments of the genome. This implies the unassembled/unaligned regions are highly enriched for high copy repeats too complex to be assembled. In contrast, the genic regions are very well represented suggesting it would be possible for a detailed analysis of the “gene-space” of the accessions from these assemblies.

Tables 2a-2c summarize the alignments of the three *de novo* assemblies relative to the reference IRGSP-1.0 Nipponbare assembly. As expected, the IR64 and DJ123 assemblies show noticeably lower overall identity, and have considerably more unaligned bases. The average k-mer coverage of the unaligned bases indicates most regions are unassembled high copy repeats, although there are 11.8 Mbp and 12.3 Mbp unaligned reference bases in IR64 and DJ123, respectively that are not repetitive based on the k-mer analysis. This suggests there may be megabases of sequence specific to each of the three genomes.

### Whole Genome Comparison to indica reference genomes

Using the same methods used for comparing to the reference Nipponbare genome, we also evaluated the three genomes relative to the reference *indica* genome (cv. 93-11) (28) (Tables 3a-3c). The agreement between the *de novo* IR64 assembly and the reference *indica* sequence is appreciably less than the Nipponbare-Nipponbare alignment; 4.31% of the IR64 assembly does not align to the 93-11 reference and the aligned regions have only 99.52% percent identity between these two *indica* varieties. Since the assemblies and alignments were computed with the same sample preparation and analysis algorithms, this suggests there are more true biological variations between IR64 and 93-11 (as would be expected from two different varieties), and/or that the 93-11 reference assembly is not as complete nor as accurate as the reference Nipponbare assembly. The later explanation is quite likely to be a contributing factor, given the fact that the 93-11 genome represents a whole genome shotgun assembly, while the Nipponbare genome utilized a combination of BACs and WGS. For example, the 93-11 assembly has 14.1 million unresolved (“N”) bases, while the Nipponbare reference has only 118.2 thousand. As seen with Nipponbare, most of the unassembled/unaligned bases between the 93-11 reference and our assemblies are repetitive with mean k-mer coverage over 14,000x. A quarter of the unaligned references bases (7.75Mbp / 31 Mbp) are non-repetitive from the k-mer analysis, while less than 900 kbp of the unaligned reference Nipponbare genome are not repetitive. This underscores the fact that there are substantially more true biological differences between IR64 and the reference 93-11 *indica* assembly than our Nipponbare sample and reference.

**Table 3a.** Comparison of the three de novo assemblies to the Indica reference (93-11 from BGI)

|  | **Unaligned**<br>**Reference**<br>**Bases** | **Unaligned**<br>**Assembly**<br>**Bases** | **Avg ID** | **SNPs &**<br>**Smal Indels** | **Indels**<br>**>5kbp** | **Inver.** | **Reloc.** | **Trans.** |
| --- | --- | --- | --- | --- | --- | --- | --- | --- |
| Nipponbare | 16.95% | 8.90% | 98.91% | 2,813,076 | 75,944 | 1,162 | 11,627 | 11,030 |
| IR64 | 13.29% | 4.31% | 99.52% | 1,228,732 | 35,762 | 644 | 6,985 | 7,101 |
| DJ123 | 14.37% | 6.78% | 99.16% | 2,264,541 | 62,191 | 974 | 8,582 | 10,170 |

**Table 3b.** Summary of unaligned reference regions relative to the Indica reference.

|  | **total bp** | **regions** | **avg size** | **max size** | **mean**<br>**k-mer coverage** |
| --- | --- | --- | --- | --- | --- |
| Nipponbare | 46,101,370 | 176,504 | 261 +/- 796 | 53,066 | 6,945 |
| IR64 | 31,006,053 | 139,336 | 264 +/- 718 | 54,056 | 14,215 |
| DJ123 | 49,562,877 | 152,378 | 325 +/- 845 | 54,307 | 9,605 |

**Table 3c.** Summary of unaligned bases by reference annotation relative to the Indica refernece. Note that only CDS and mRNA annotations are available for the reference Indica assembly.

|  | **mRNA** | **CDS** | **repeat bp (>100x**<br>**kmer coverage)** |
| --- | --- | --- | --- |
| Nipponbare | 9,768,022 | 4,809,510 | 33,934,310 |
| IR64 | 7,629,670 | 3,993,153 | 23,255,924 |
| DJ123 | 5,999,753 | 1,947,112 | 23,805,398 |

Finally, the comparison between the IR64 and the DJ123 assemblies shows that they differ from each other nearly as much as either one differs from the Nipponbare reference sequence. These results suggest that the *aus* genome harbors a greater amount of novel variation than previously recognized. It also highlights the value of taking an unbiased, *de novo* assembly approach when evaluating genomic variation among varieties and subpopulations to capture genome-specific variations.

### Pan-genome analysis

We next evaluated the “pan-genome” of the three *de novo* assemblies to identify sequences that were conserved across the genomes as well as sequences specific to just one genome (see methods). Using the whole genome alignment information, we classified each base of each genome as being specific to that genome (unaligned to either other genome), or shared by one or both genomes. The majority of the assembled sequences (∼302 Mbp per genome) and exonic sequences (∼55.5Mbp per genome), were shared among the three genomes, although 4.8 Mbp to 8.2 Mbp (423 kbp to 930 kbp exonic) were found to be genome-specific (Figure 2a). Since a gene sequence may be partially shared or partially genome-specific, we assigned each gene to the sector on the Venn diagram for which the majority of the exonic bases were assigned over all transcripts associated with each gene. For example if 90% of a gene is shared among all three genomes, but 10% is genome-specific, we would assign it to the center (fully shared) sector under the majority rule. This will not necessarily characterize changes in gene function if critical protein domains are shared/unshared, but highlights the major trends between the lineages and discovers 297 to 786 genome-specific loci.

**Figure 2.**
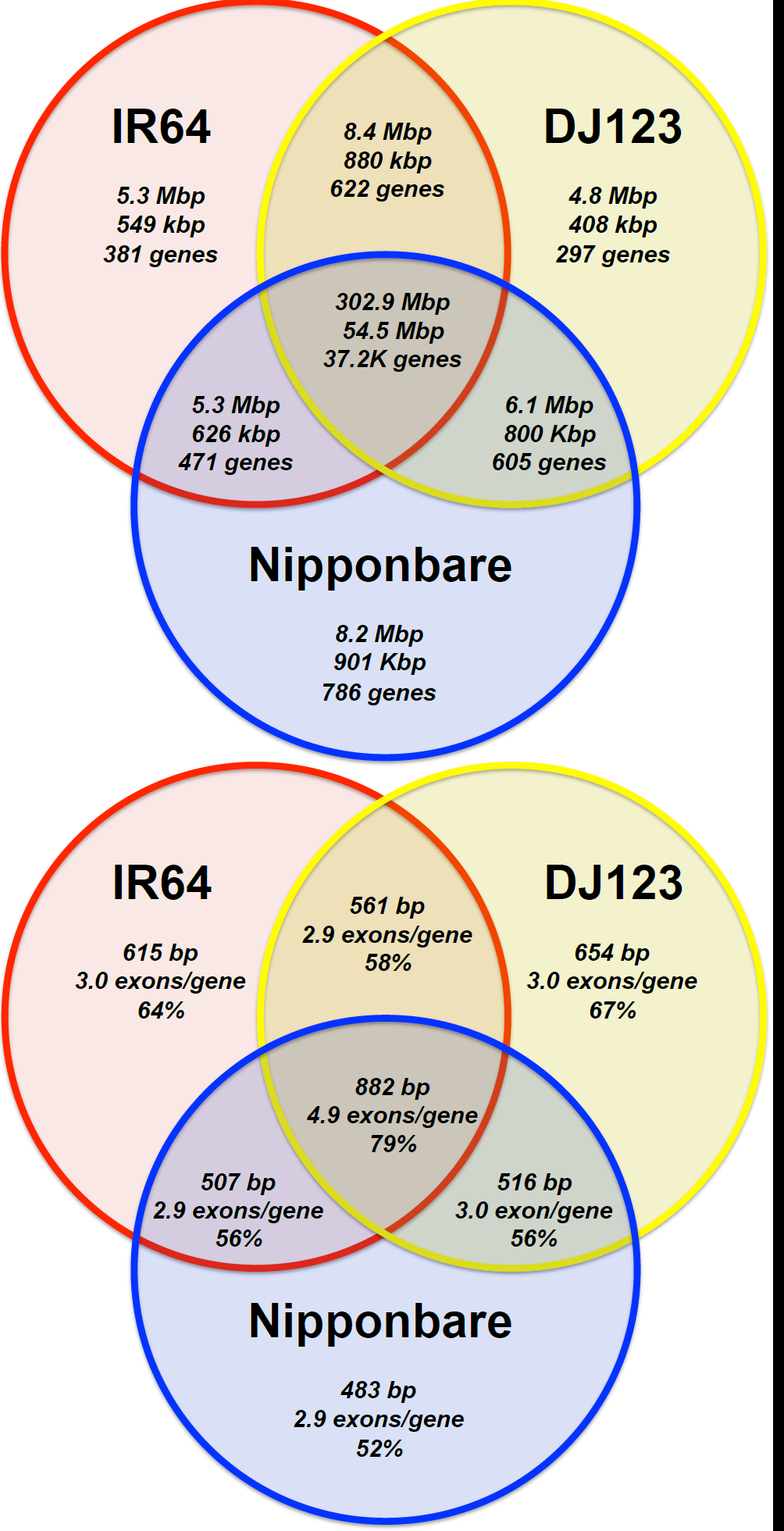
Venn diagrams of the shared sequence content between Nipponbare (*temperate japonica)*, IR64 (*indica)* and DJ123 (*aus*) **(A)** overall sequence content. In each sector, the top number is the total number of base pairs, the middle number is the number of exonic bases, and the bottom is the gene count. If a gene is partially shared, it is assigned to the sector with the most exonic bases. **(B)** genic content. In each sector, the top number is the median CDS length, the middle number is the average number of exons/gene, and the bottom is the percentage InterPro/homology.

Using the same k-mer analysis techniques we applied for the reference analysis, we further classified the genome-specific bases as being unique or repetitive, using a threshold of 100x average k-mer coverage to classify unique sequences. From this, we identified only 1.2 Mbp to 1.5 Mbp of nonrepetitive sequence specific to each genome, meaning that most of the genome-specific bases were actually repetitive (Table 4). Since repetitive sequences are also the most likely to be unassembled, as observed in our comparison to the reference genomes, we further examined the genome-specific exonic bases and refined our initial estimates to 555 kbp to 760 kbp of non-repetitive, genome-specific sequences intersecting annotated genes by at least 100 bp (Table 5). Note these segments may include flanking promoter and other regulatory regions in addition to the exons themselves. From this catalog, we selected 10 of the largest regions in each of the genomes for PCR validation, and were able to confirm the computational analysis with 100% success (Figure 3, Supplementary Table 4a-4c).

**Table 4.** Genome-specific non-repetitive bases. Identified sequences must be at least 100 bp long with no alignments to the other genomes, average between 10x and 100x kmer coverage in that genome, and average below 10x kmer coverage using the reads from the other two genomes.

| Genome | Genome-specific | Regions | mean +/- stdev | max size |
| --- | --- | --- | --- | --- |
| Nipponbare | 1,574,801 bp | 2,250 | 699 +/- 904 | 8,463 |
| IR64 | 1,336,650 bp | 1,702 | 785 +/- 1066 | 10,400 |
| DJ123 | 1,263,681 bp | 1,569 | 805 +/- 1058 | 8,960 |

**Table 5.** Genome-specific non-repetitive gene sequences. Regions identified to be specific to a given accession using the criterion used in Table 4 and intersect an annotated gene region by at least 100 bp.

| Genome | Genome-specific | Regions | mean +/- stdev | max size |
| --- | --- | --- | --- | --- |
| Nipponbare | 760,064 bp | 779 | 975 +/- 1,197 | 8,463 |
| IR64 | 637,470 bp | 583 | 1,093 +/- 1,430 | 10,400 |
| DJ123 | 555,507 bp | 492 | 1,129 +/- 1,409 | 8,960 |

**Figure 3.**
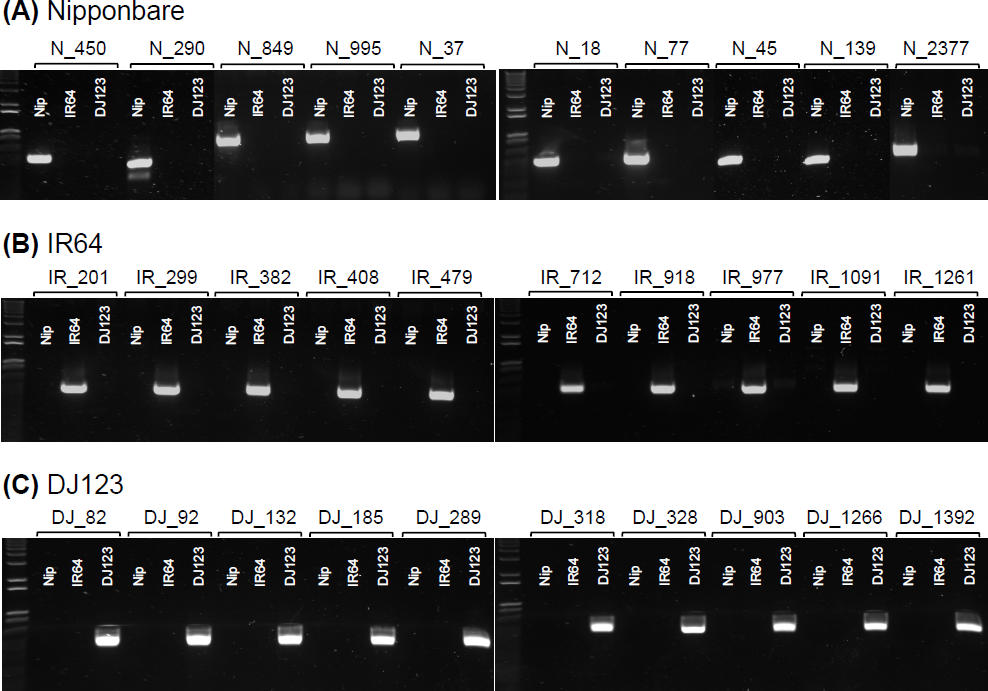
PCR validation of genome-specific regions. Regions identified as unique to each genome assembly were amplified from genomic DNA of all three genomes and visualized on 1% agarose gels. (**A**) Nipponbare-specific sequences; (**B**) IR64-specific sequences, (**C**) DJ123-specific sequences.

For Nipponbare and IR64, we determined the positions of the non-repetitive segments along the different reference chromosomes, and found the segments were broadly distributed. For Nipponbare, we could localize 2,208 of the genome-specific regions, and found that one region occurred, on average, every 162 kbp +/− 362 kbp, following an approximately exponential distribution (data not shown). For IR64, we could localize 1,074 of the genome specific-regions, and found one region occurred on average every 338 kbp +/− 752 kbp, also from an approximately exponential distribution. The distributions suggest that the genome-specific bases are not highly localized, as an exponential distribution in spacing can occur if there is a uniform probability distribution of a site occurring at any position at random.

Genome-specific loci, as well as those shared between two genomes but not the third, exhibited shorter coding-sequences and greater novelty compared to genes shared among all three genomes (Figure 2b). For example, loci common to all genomes had a median coding-length of 888 bp compared to median values ranging from 483-654 bp for the genome-specific gene sets. Likewise, the core, fully shared set of genes averaged 4.9 exons/transcript compared to a range of 2.9-3.0 genes/transcript amongst genome-specific genes. A smaller fraction of genome-specific loci contained InterPro domains compared to the core set (40% vs 63%), and fewer showed homology to plant RefSeq proteins (57% vs 79%). Similar to these findings, studies in yeast, Drosophila, and vertebrates have found that novel and recently evolved genes tend to encode smaller proteins than conserved or ancient genes (52–54). However, artifacts of inaccurate annotation may also contribute to this trend (54).

To characterize potential function of genome-specific genes we further examined genes with annotated InterPro domains. Notably, genes with domains related to disease resistance were the most prevalent type among genome-specific genes. For example, 12% of genes specific to IR64 possessed the NB-ARC motif (IPR002182), the central nucleotide-binding domain of plant R-genes. This domain, and others associated with R-genes, also prevailed among the DJ123-specific and Nipponbare-specific gene sets, accounting for 9% and 5% of genes respectively. In contrast, only 0.35% of genes shared among all three genomes encode the NB-ARC domain. Genes shared between just two genomes showed intermediate frequencies of disease resistance genes (1.5 to 2.5 %). Similar distributions were seen for genes classified with the gene ontology term “defense response” (GO:0006952). These results are consistent with Ding et al. (2007), who showed high levels of “genome asymmetry” among R genes when comparing the Nipponbare and 93-11 reference assemblies (13). A large diversity of other protein-domain classes, such as those associated with receptor and non-receptor protein kinases, transcription factors, metabolic enzymes, proteases, and transporters were also found in the genome-specific gene sets. A complete listing of putative strain-specific genes, their InterPro domains, GO terms, and summary of homology search results, are provided in a supplementary file.

### Detailed regions

We chose four agronomically relevant regions of the rice genome that were previously reported to harbor differences among the three varieties or subpopulations to illustrate the utility of these high quality whole genome assemblies for understanding the variation in genome structure underlying salient phenotypic variants.

#### 1. *S5* hybrid sterility locus

*S5* is a major locus for hybrid sterility in rice that affects embryo-sac fertility. Genetic analysis of the *S5* locus documented three alleles: an *indica* (*S5*-i), a *japonica* (*S5*-j), and a neutral allele (*S5*-n) (23, 55). Hybrids of genotype *S5*-i/*S5*-j are mostly sterile, whereas hybrids of genotypes consisting of *S5*-n with either *S5*-i or *S5*-j are mostly fertile. The *S5* locus contains three tightly linked genes that work together in a ‘killer-protector’ type-system (56, 57). During female sporogenesis, ORF5+ (killer) and ORF4+ (partner) cause endoplasmic reticulum (ER) stress. ORF3+ prevents ER stress and allows the production of normal gametes, whereas the ORF3- allele cannot prevent it, resulting in embryo-sac abortion. The *ORF3-* allele has a 13-bp deletion; the *ORF4-* allele carries an 11-bp deletion that causes a premature stop codon (57). The ORF5 *indica (ORF5+)* and *japonica (ORF5-)* alleles differ by only two nucleotides, whereas the wide compatibility allele *S5-n (ORF5n)* has a large deletion in the N-terminus of the predicted protein, rendering it presumably nonfunctional (56). The typical *indica* haplotype is ORF3+/ORF4-/ORF5+, while the typical *japonica* haplotype is ORF3−/ORF4+/ORF5−.

In each of the three *de novo* assemblies reported here the *S5* locus containing the three genes lies within a single scaffold and haplotypes can be easily identified (Supplementary Table 5, Supplementary Figure 2). The identity of the *ORF5* alleles in Nipponbare, IR64 and DJ123 were also confirmed by Sanger sequencing from genomic DNA, and perfectly confirm the assembly results (Supplementary Figures 3-4). The Nipponbare assembly is in agreement with the Nipponbare IRGSP-1.0 reference sequence for the region and shows that it carries the typical *japonica* haplotype ORF3−/ORF4+/ORF5−. The IR64 assembly shows that this accession carries the typical *indica* haplotype ORF3+/ORF4−/ORF5+. In the case of DJ123, our *de novo* assembly revealed that this *aus* accession carries the 136-bp deletion characteristic of the neutral allele, *ORF5n.* However, the DJ123 *ORF5n* allele is novel, as it differs from the reported *ORF5n* allele by two SNPs and one 10-bp deletion within the coding region of the gene (also confirmed by Sanger sequencing). The DJ123 haplotype for the locus is ORF3−/ORF4−/ORF5n, a haplotype previously identified by Yang *et al.* (2012) in four accessions from Bangladesh. Although the accessions bearing this haplotype were referred to as *indica* in this study (57), they almost certainly belonged to the *aus* subpopulation.

#### 2. *Sub1* locus

The *Submergence 1 (Sub1)* locus on chromosome 9 is a major QTL determining submergence tolerance in rice (33). The *Sub1* locus is a cluster of three genes encoding putative ethylene response factors. *Sub1B* and *Sub1C* are present in all rice accessions tested to date, while *Sub1A* may be present or absent. Originally identified in the *aus* accession FR13A, *Sub1A* appears to be found only within the *Indica* varietal group (33). *Sub1A* has two alleles: *Sub1A-1* is found in submergence-tolerant varieties, while *Sub1A-2* is found in intolerant varieties. A haplotype survey in *O. sativa* varieties also identified nine *Sub1B* and seven *Sub1C* alleles (33).

In the IR64 and DJ123 *de novo* assemblies reported here the *Sub1* locus lies within a single scaffold and haplotypes can be easily identified (Supplementary Table 5). In the IR64 assembly the *Sub1A* gene is present as the *Sub1A-2* allele, previously identified in submergence-intolerant accessions including IR64 (33). For the *Sub1B* and *C* genes, IR64 carries the alleles *Sub1B-1* and *Sub1C-3*, as reported (33). *Sub1A* is absent from the DJ123 assembly, suggesting that this *aus* variety is not submergence tolerant. DJ123 carries a novel *Sub1B* allele (Sub1B-10), and the previously identified *Sub1C-6* allele. In the Nipponbare assembly, *Sub1B* and *Sub1C* lie within a single scaffold, and the alleles identified are in agreement with published results (33). Nipponbare is not submergence tolerant and the *Sub1A* gene is absent in Nipponbare according to previous reports. Our *de novo* assembly is unresolved in the region that corresponds to the *Sub1A* gene, but a k-mer analysis using the methods and data applied above clearly shows a lack of coverage in the DJ123 and Nipponbare sequencing reads across the locus except for high copy repeats dispersed in the sequence (Fig. 4 top and bottom). Conversely, the k-mer coverage of the IR64 assembly is uniformly at the single-copy coverage level (∼100x), except for a small number of localized gaps in coverage, corresponding to SNPs in IR64 relative to the reference *Sub1A* sequence, and the high frequency repeats (Fig. 4 middle). In contrast, the k-mer coverage across *Sub1B* and *Sub1C* is consistently ∼100x, except for isolated sharp gaps corresponding to variations relative to the reference sequences (Supplementary Figs. 5 - 6).

**Figure 4.**
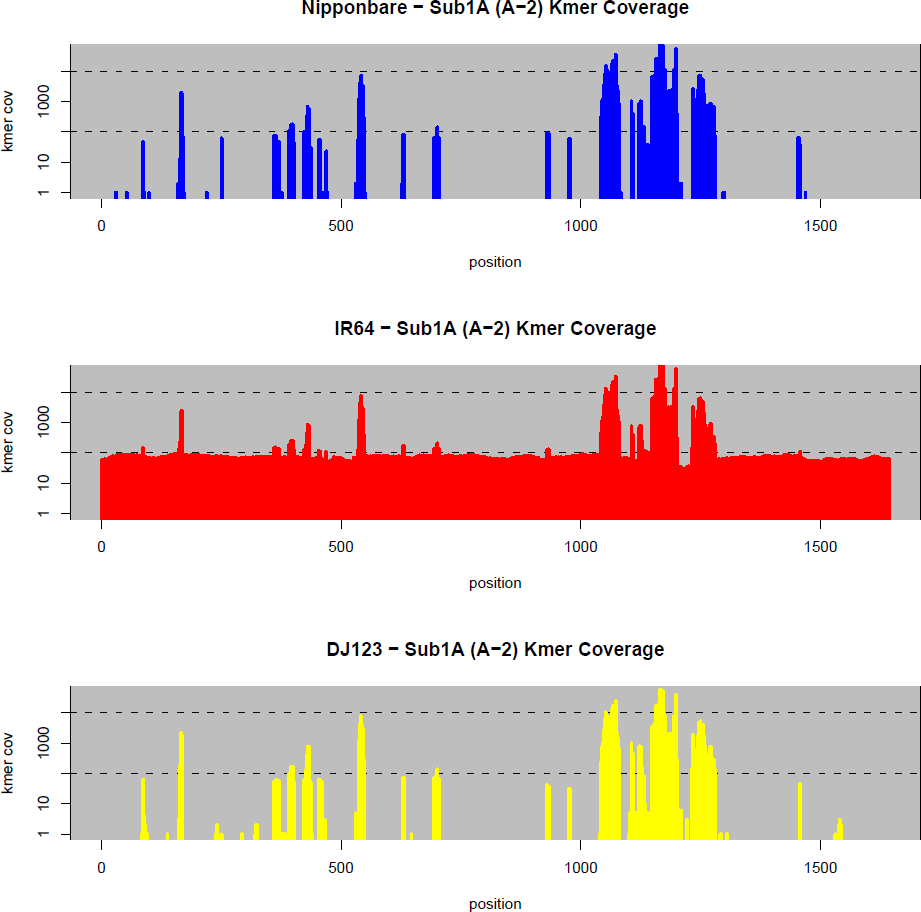
K-mer coverage in the three assemblies across the *Sub1A* gene. In each panel, the k-mer coverage of the sequence reads of the 3 respective genomes are plotted according to the sequence of the Sub1A A-2 allele. Only IR64 has consistent coverage across the gene, while the other two genomes have sparse coverage of a few repetitive k-mer sequences. For clarity, the k-mer coverage range 1x to 50,000x (log scale) is displayed in all the plots.

#### 3. *LRK* gene cluster

Fine-mapping of a yield-improving QTL on rice chromosome 2 identified a cluster of leucine-rich repeat receptor kinase genes (58), consisting of seven or eight intronless gene copies contained within a 40 – 50 Kb genomic region. The QTL, originally introgressed from a wild rice accession (Dongxiang), was shown to increase grain yield of the recurrent parent Guichao2 (*indica)* by about 25%. The *LRK* locus in Dongxiang carries an extra gene, *LRK1*, absent from Guichao2. A survey of haplotype divergence in 13 rice accessions showed that *LRK1* is absent in only three *indica* accessions, suggesting that these haplotypes may have originated via gene loss.

In each of the three *de novo* assemblies reported here the *LRK* locus lies within a single scaffold and haplotypes can be easily identified (Supplementary Table 5). The Nipponbare assembly is in agreement with the reference sequence, with the exception of regions that the *de novo* assembly was not able to resolve because of high copy repeats (Supplementary Figure 7). *LRK1* is absent in the IR64 assembly as evident in the k-mer plot (Supplementary Figure 8), indicating that IR64 carries the 7-gene haplotype identified in other *indica* accessions (58). According to our assembly and the corresponding k-mer analysis, the *aus* accession DJ123 carries *LRK1.* Based on sequence variation on the 5’ upstream region of *LRK4* and *LRK6*, we can predict that the DJ123 haplotype for the LRK gene cluster is closest to the haplotypes identified in *indica* accessions in which *LRK1* is present (haplotypes A, B and C in Fig. 3 of (58)).

#### 4. *Pup1* region

*Phosphorus uptake1 (Pup1)* is a major rice QTL associated with tolerance to phosphorus deficiency in soils (59, 60). The *Pup1* locus is a large, 90 Kb region originally identified in Kasalath, an *aus* variety that is tolerant to phosphorus deficiency, but is absent in phosphorus starvation-intolerant varieties, including Nipponbare (61). A gene encoding a protein kinase, *Pstol1*, located within the 90 Kb indel, is responsible for the P-uptake efficiency phenotype (40).

Of the three *de novo* assemblies reported here, the 90 Kb indel is absent from both Nipponbare and IR64, but a large portion of it, including the *Pstol1* gene, is present in the *aus* variety DJ123 (Figure 5, Supplementary Table 5). Although it is at least partially present, the region of the 90 Kb indel described in Kasalath could not be fully resolved in our DJ123 assembly. This suggests that the 90 Kb indel may be truncated and/or rearranged in some *aus* varieties. Interestingly, as shown in Fig. 5, the Kasalath reference sequence contains unresolved gaps flanking regions of very high k-mer coverage, therefore longer reads may be necessary to assemble this region with confidence. The *Pstol1* gene sequence is complete in DJ123, and shows six SNPs relative to the Kasalath sequence (also apparent as abrupt drops in coverage in the k-mer coverage plot, Supplementary Fig. 9-10). These SNPs were confirmed via Sanger sequencing on genomic DNA (Supplementary Fig. 11). One of these SNPs introduces a premature stop codon, resulting a protein that is only 136-amino acids long (the intact PSTOL1 protein is 324 aa), therefore presumably non-functional.

**Figure 5.**
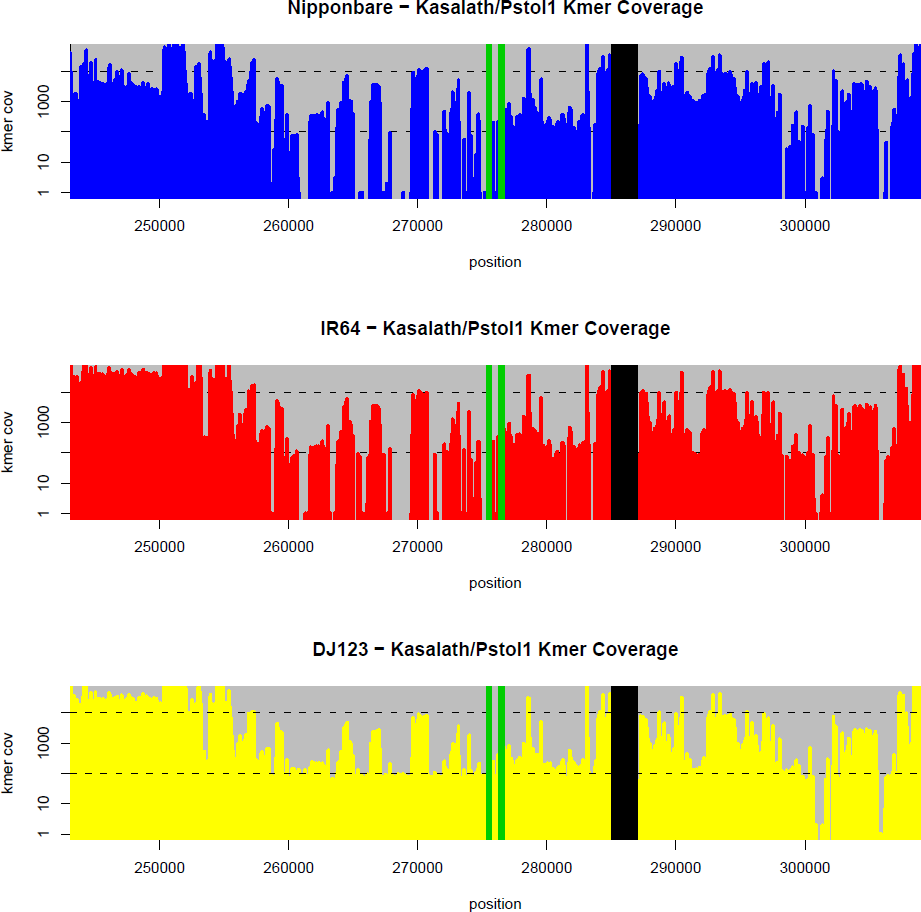
K-mer Coverage across the Kasalath*/Pstol1* gene in the three genomes with 30 kbp of upstream and downstream flanking sequence. The k-mer coverage is plotted with respect to the reference Kasalath sequence (AB458444.1). The position of the *Pstol1* gene is indicated with green vertical bars. Also see Supplementary Figure 9 for detailed view of the *Pstol1* coverage, and Supplementary Figure 10 for a plot of the entire Kasalath sequence. Unresolved gaps in the reference sequence are indicated with black vertical bars. Only DJ123 has consistent coverage across this region, especially upstream of the gene, while the other two genomes show complete gaps in coverage.

## Discussion

In this study we wanted to overcome the limitation on sequencing and comparison to a reference genome by developing high quality *de novo* assemblies to observe biologically significant changes between multiple rice genomes. The rice accessions sequenced were selected to represent the *indica* (cv IR64), *aus* (DJ123) and *temperate japonica* (Nipponbare) subpopulations (**Figure 1**). The inclusion of the high quality, BAC-by-BAC assembly of Nipponbare and the shotgun assembly of 93-11 provided a control that allowed us to assess the quality of the different datasets and *de novo* assembly strategies. It is apparent from comparing different assembly software that ALLPATHS-LG gave the best results in our hands (Supplemental Table 3). It is also apparent that the use of k-mer frequencies is a robust technique for characterizing repetitive regions, and enabled us to correctly characterize and validate genome-specific regions.

The three-way comparison among the different genomes was informative in identifying major shared and structurally variable regions of the rice genome. We were particularly interested in regions that were structurally unique to either the *indica* and/or the *aus* genome because they would likely have been discarded in previous re-sequencing efforts due to difficulties aligning their sequencing reads to the Nipponbare reference genome. This would be particularly true for longer genome-specific sequences, which would be completely absent in the alignments to the reference. We anticipate future studies will systematically perform follow up functional studies of these loci as being likely candidates for phenotypic differences observed between the genomes.

Our analysis clearly demonstrates that the *indica* and the *aus* genomes are more distantly related than previously known. Because the *aus* subpopulation is phenotypically so similar to *indica*, the degree of genetic differentiation has been underappreciated by breeders and geneticists alike (42, 62, 63). The unusual characteristics of the *aus* subpopulation, combined with evidence of unique *aus* alleles at loci such as *Rc*, conferring white versus colored pericarp (18), the *Snorkel* locus conferring deep water ability (44), the *Pstol1* locus conferring phosphorus-update efficiency (40), or the *Sub1* locus conferring submergence tolerance (33), all support the hypothesis that *aus* may have a unique domestication history compared to *japonica* and *indica*. These findings underscore the importance of recognizing genetic subpopulation structure to guide plant breeders in identifying novel sources of variation for traits of interest. In recent years, many key biotic and abiotic stress tolerance genes have been discovered in *aus* varieties (33, 40, 43–45). It is interesting to note that in several cases, the donor *aus* germplasm is referred to as *indica*, underscoring how *indica* and *aus* are often confused, as noted for the DJ123 haplotype of the S5 hybrid sterility locus (see above).

The overall annotation of our Nipponbare assembly is quite close to that of the reference Nipponbare genome. This illustrates that the approach we describe here provides a genome sequence of considerable vitality for further research. However, our contig N50 sizes (as opposed to scaffold N50) are still fragmented by the presence of complex repeats, which somewhat limits application when studying large structural rearrangements, as exemplified by the Pup1 region, which remains partially unassembled in the *aus* variety DJ123 (Fig. 6). We anticipate that some combination of short-read NGS sequencing and newly emerging long read sequences, such as PacBio Single Molecule Real Time Sequence (SMRT) (64), will soon overcome this limitation and provide assemblies approaching, or perhaps even surpassing, those provided by the vastly more expensive and time consuming BAC-by-BAC approach. Once this occurs it should spark an outburst of genomics studies of agronomically important plant genomes, greatly enriching our potential to understand their many unique qualities and characteristics and paving the way for enhanced utilization of natural variation in plant improvement.

## Methods

### Plant material

Three rice (*Oryza sativa)* accessions (Nipponbare, IR64, DJ123) were used in the study. Accession information (i.e. Genetic Stocks *Oryza (GSOR)* identifier, accession name, country of origin, subpopulation) is summarized in Table 6 (62). The plants were grown in the Guterman greenhouse facility at Cornell University, leaf tissue was harvested from one-month-old seedlings, ground in a mortar and pestle, and DNA was extracted using the Qiagen Plant DNeasy kit (Qiagen, Valencia, CA, USA).

**Table 6.**
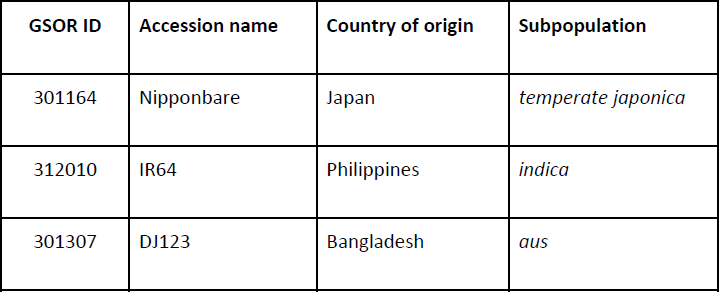
Accession information for the three rice genomes in the Genetic Stocks *Oryza* (GSOR) stock center.

| **GSOR ID** | **Accession name** | **Country of origin** | **Subpopulation** |
| --- | --- | --- | --- |
| 301164 | Nipponbare | Japan | *temperate japonica* |
| 312010 | IR64 | Philippines | *indica* |
| 301307 | DJ123 | Bangladesh | *aus* |

### DNA sequencing

The DNA sequencing was performed in the Cold Spring Harbor Laboratory Genome Center using Illumina HiSeq 2000 instruments. For each of the three varieties, three libraries were sequenced following the requirements and recommendations of the ALLPATHS-LG whole genome assembler: (1) a 180 bp fragment library sequenced as 2x100 bp reads; (2) a ∼2 kbp jumping library sequenced as 2x50 bp reads; and (3) a ∼5 kbp jumping library sequenced as 2x50 bp reads.

For the 180 bp overlap library the sample was mechanically fragmented by using the Covaris S2 System and then prepared based on the New England Biolabs NEBNext Illumina library protocol and ligated to standard Illumina paired-end adapters. To maximize sample throughput the samples were size-selected in 50-bp windows between 290 - 310 bp using the Caliper XT instrument. Each library was PCR enriched for 12 cycles and quantified using the Bioanalyzer.

For the jumping libraries, the Illumina mate-pair library protocol was used. The DNA was fragmented into 2 kb and 5 kb segments. We again used the Covaris S2 System using programs that we developed in the lab. The fragmented DNA was then end-repaired with biotin-labeled dNTPs. The labeled fragments are circularized and fragmented again into 400 bp pieces. Fragments with the biotin labels are enriched, end-repaired, and ligated with adapters used for downstream processes. Each library was PCR enriched for 18 cycles and size-selected for 350-650 bp fragments. The final library consists of fragments made up of two DNA segments that were originally separated by ∼2 kbp or ∼5kb. Each of the libraries was sequenced to 30x-80x sequence coverage, as recommended by the assembler.

Libraries were sequenced on one or more lanes of an Illumina HiSeq 2000 using paired-end 50 or 100 bp runs. Image processing and basecalling were performed as the runs progressed with Illumina’s Real Time Analysis (RTA) software. The binary basecall files were streamed to a shared Linux server for further processing. The Illumina Casava pipeline (v1.8) was used to process the binary files to fastq files containing the basecalled reads and per base quality scores. Only reads passing the standard Illumina quality filter were included in the output files.

### Genome Assembly

The ALLPATHS-LG version R41348 assembly algorithm was used for the assemblies. It consists of 5 major phases: (1) pre-assembly error correction, (2) merging of the overlapping fragment reads into extended reads, (3) constructing the unipath graph from the k-mers present in the reads, (4) scaffolding the unipaths with the jumping libraries, and (5) gap closing. To complete the 5 phases, the algorithm requires an overlapping pair fragment library and at least one jumping library, although the authors recommend at least 2 jumping libraries of ∼2 kbp and ∼5 kbp or larger. We assembled each of the genomes using ∼50x coverage of the fragment library and ∼30x coverage of each of the two jumping libraries using the recommended parameters, except we lowered the MIN_CONTIG size to 300 bp from the default 1000 bp. This parameter controls the minimum contig size to be used for scaffolding, and our previous testing determined this change leads to (modestly) improved contig and scaffold statistics.

We also evaluated using SOAPdenovo2 (65) and SGA (66) for the assemblies (Supplementary Table 3), using the same fragment, 2 kbp, and 5 kbp libraries but both assemblers had substantially worse contiguity statistics under a variety of parameter settings. For SOAPdenovo2, we corrected the reads using the Quake error correction algorithm (67), and then ran 7 assemblies with the de Bruijn graph kmer size set to k=31 through k=45 (odd values only, as required). In every attempt the scaffold N50 size was below 10 kbp compared to >200 kbp for our best ALLPATHS-LG assembly. For SGA, we evaluated 4 assemblies with the string graph minimum overlap length of k=71 through k=77 (odd values only, as required), but the scaffold N50 size was below 15 kbp in every attempt. We hypothesize that ALLPATHS-LG achieved superior results because the algorithm automatically measures many of the properties of the sequencing data, and could therefore self-adjust the various cutoffs used by the algorithm for error correction, contigging, and scaffolding.

Applying nomenclature proposed by (50), we have named these assemblies to convey accession, quality, origin, and iteration as follows: Os-Nipponbare-Draft-CSHL-1.0, Os-IR64-Draft-CSHL-1.0, Os-DJ123-Draft-CSHL-1.0.

### Genome Annotation

Repeat elements were masked using RepeatMasker (68) with a rice repeat library available from the Arizona Genome Institute (AGI). Protein-coding genes were annotated using MAKER-P version 2.30, installed on the Texas Advanced Computer Center Lonestar cluster and provisioned through an iPlant Collaborative allocation (51, 69–71). Sequence evidence used as input for MAKER-P included *Oryza* expressed sequences (EST, cDNA, and mRNA) downloaded from the National Center for Biotechnology Information (NCBI), and annotated coding and protein sequences available for Nipponbare (IRGSP1.0 and MSU release 7) (50), 93-11 (28), and PA64s (Supplementary Table 1). *Ab initio* gene predictions made using FGENESH (72) were incorporated exogenously into the MAKER-P pipeline using the pred_gff parameter. The SNAP (73) *ab initio* predictor was run within MAKER-P using the O.sativa.hmm parameter provided with SNAP. To annotate protein domain structure and assign Gene Ontology (GO) terms we used InterProScan 5 software (74), available within the iPlant Discovery Environment (75). Among resulting InterPro domains we curated 21 as being associated with transposon-encoded genes and screened out MAKER-P annotations with these domains (IPR000477,IPR001207,IPR001584,IPR002559,IPR004242,IPR004252,IPR004264,IPR004330,IPR0043 32,IPR005063,IPR005162,IPR006912,IPR007321,IPR013103,IPR013242,IPR014736,IPR015401,IPR01 8289,IPR026103,IPR026960,IPR027806). To identify homologies we conducted BLASTP alignment to the plants sub-section of NCBI RefSeq (release 63), using an e-value threshold of 1e-10.

### Whole Genome Comparisons

We used the MUMmer (76) whole genome alignment package and the GAGE assembly comparison scripts to compare the *de novo* assemblies to the reference Nipponbare and *Indica* genomes. Briefly, we aligned the assemblies to the genomes using *nucmer* using sensitive alignment settings (−c 65 −l 30 -banded −D 5). For base level accuracy evaluations, we used the GAGE assembly comparison script, which further refines the alignments by computing the best set of 1-to-1 alignments between the two genomes using the dynamic programming algorithm *delta-filter.* This algorithm weighs the length of the alignments and their percent identity to select 1-to-1 non-redundant alignments. This effectively discards spurious repetitive alignments from consideration, allowing us to focus on the meaningful differences between the genomes. Finally, the evaluation algorithm uses *dnadiff* to scan the remaining, non-repetitive alignments to summarize the agreement between the sequences, including characterizing the nature of any non-aligning bases as substitutions, small indels, or other larger structural variations. To characterize the unaligned regions of the reference genome, we converted the whole genome alignments into BED format. For this we did not exclude repetitive alignments, so that we could focus on novel sequence instead of copy number differences. We used BEDTools (77) to intersect the unaligned segments with the reference annotation, and summarized the size distributions of the unaligned segments using AMOS (78).

#### K-mer Analysis

To evaluate the repeat composition, we selected a random sample of 400M unassembled reads from each of the three genomes and used *Jellyfish* (79) to count the number of occurrences of all length 21 k-mers in each read set. Length 21 was selected to be sufficiently long so that the expected number of occurrences of a random k-mer was below 1, but short enough to be robust to sequencing errors. The modes of the 3 kmer frequency distributions, excluding erroneous k-mers that occurred less than 10 times, were 60x (Nipponbare), 64x (DJ123), and 73x (IR64) drawn from an approximately negative binomial distribution (Supplementary Figure 1). These values correspond to the average k-mer coverage for single copy, non-repetitive regions of the genome. See Kelly et al. (67) for a discussion of k-mer frequencies. We then used the AMOS program *kmer-cov-plot* (78) to report the kmer coverage along the two reference genomes using the 3 databases of read k-mer frequencies. Unlike read alignments, which may be sensitive to repeats and variations, evaluating k-mer coverage is very robust to determine repetitive content (80, 81). Single nucleotide variants are also readily apparent in these plots as abrupt gaps in coverage k-bp long, while indels will be present as longer gaps in coverage (82).

#### Pan-Genome Analysis

The pan-genome analysis followed the reference-based analysis above, using *nucmer* to align the genomes to each other, *BEDTools* to find the genome-specific and shared regions of the genomes, and the *jellyfish/AMOS* k-mer analysis as described above to classify unique and repetitive sequences. We also used *BEDTools* to intersect the genome-specific/shared regions against their respective annotations to determine how the exonic bases were shared across the genomes. We summarized the genome-specific/shared exonic bases into gene counts by counting the total number of shared or specific exonic bases across all possible transcripts for a gene, and assigned the gene to the sector of the Venn diagram with the most bases associated with it. For the purposes of the Venn diagram (Fig. 2a), wherever possible, the Nipponbare base or gene counts were used, followed by the values from IR64, and then followed by the DJ123 specific values, although the values were all largely consistent.

#### PCR and sequencing validation of specific regions

The same algorithms and parameters as the Pan-Genome analysis were also used to characterize the specific regions identified in the paper. PCR and/or sequencing validation were performed on genomic DNA extracted from tissue collected from independently grown plants obtained from the same seed source used for Illumina sequencing. Genomic DNA was extracted from young leaf tissue using the Qiagen Plant DNeasy Mini kit. Primers used for validation of 10 of the longest genome-specific sequences from each rice line, and of the *S5* and *Pup1* loci, are listed in Supplementary Table 4a-4c. Sanger sequencing was performed at the Biotechnology Resource Center at Cornell University.

## Data Access

The read data, assemblies, annotations, and pan-genome alignments are posted on the CSHL website at http://schatzlab.cshl.edu/data/rice, including the accession numbers for the short read data in the NCBI SRA. Analysis software packages are available open source from the websites for ALLPATHS-LG (http://www.broadinstitute.org/software/allpaths-lg/blog/?pageid=12), MUMmer (http://mummer.sourceforge.net), AMOS (http://amos.sourceforge.net), Jellyfish(http://www.genome.umd.edu/jellyfish.html), and BEDTools (https://github.com/arq5x/bedtools2).

## Acknowledgements

This project was supported in part by National Science Foundation awards PGRP-1026555 to SMc, DBI-126383 to DW and MCS, and IOS-1032105 to WRM and DW, DBI-0933128 to WRM. It was also supported in part by National Institutes of Health award R01-HG006677 to MCS. We would like to thank Adam Phillippy and Sergey Koren for their helpful discussions with the GAGE assembly validation software and pan-genome alignments; Aaron Quinlan for his helpful discussions with BEDTools; and David Jaffe, Iain MacCallum, Ted Sharpe, Filipe Joao Ribeiro, and all the ALLPATHS-LG developers and support staff for the assistance debugging and troubleshooting the assemblies.

